# Differential Responses of Lung and Intestinal Microbiota to SARS-CoV-2 Infection: A Comparative Study of the Wuhan and Omicron Strains in K18-hACE2 tg Mice

**DOI:** 10.1101/2024.10.25.620241

**Authors:** Chae Won Kim, Keun Bon Ku, Insu Hwang, Hi Eun Jung, Kyun-Do Kim, Heung Kyu Lee

## Abstract

The COVID-19 pandemic, caused by SARS-CoV-2, has led to the emergence of viral variants with distinct characteristics. We investigated the differential effects of the original Wuhan strain and the emergent Omicron variant of SARS-CoV-2 using a K18-hACE2 transgenic mouse model. We compared the mortality rates, viral loads, and histopathological changes in lung and tracheal tissues, as well as alterations in the lung and intestinal microbiota following infection. We observed significant differences in disease severity, with the Wuhan strain causing higher mortality and more severe lung damage than the Omicron variant. Furthermore, microbiome analyses revealed distinct shifts in microbiota associated with infection by each variant, suggesting that microbiome-related mechanisms might influence disease outcomes. This comprehensive comparison enhances our understanding of COVID-19 pathogenesis and highlights the importance of microbiome dynamics in viral infections, providing insights for future therapeutic and preventive strategies.

**Importance:** Understanding the differential impacts of SARS-CoV-2 variants is crucial for effective public health response and treatment development. This study provides insights into the pathogenesis of the original Wuhan strain and the Omicron variant of SARS-CoV-2, revealing significant differences in host mortality, viral load, and lung pathology. The use of the K18-hACE2 transgenic mouse model enables detailed examination of these differences in a controlled setting. Furthermore, this study highlights the importance of the microbiome in modulating disease severity and host responses to viral infections. By uncovering distinct microbial shifts associated with infection by different SARS-CoV-2 variants, this study suggests potential microbiome-related mechanisms that might be targeted to mitigate disease outcomes.

## Introduction

SARS-CoV-2 has led to an unprecedented global health crisis, with various strains exhibiting varying levels of virulence and transmission (1, 2). Since the initial outbreak in Wuhan, China, the virus has evolved, resulting in the emergence of variants with distinct characteristics and health impacts (3–5). Among these, the original Wuhan strain (SARS-CoV-2 Wuhan) and the emergent Omicron variant (SARS-CoV-2 Omicron) are particularly notable for their differences in virulence, transmissibility, and clinical outcomes (6). The Wuhan strain, which is known for its high mortality rate, causes severe respiratory and systemic complications (7, 8). In contrast, the Omicron variant, despite its higher transmissibility, is often associated with milder symptoms and lower mortality rates (6, 9).

Understanding the differential effects of these strains on disease outcomes and host microbiota is critical for developing public health strategies and interventions (10). Changes in gut microbiota have been suggested to influence lung diseases (11–13), and this topic has garnered increased interest in the context of SARS-CoV-2 infection (14–16). As a result, SARS-CoV-2 was found to infect and affect the gut tissues that have the virus-specific entry receptor angiotensin-converting enzyme 2 (ACE2) (17, 18). SARS-CoV-2 infection was also found to be accompanied by dysbiosis of respiratory tract microbiota (19); however, there are practical limitations to analyzing lung microbiota in human patients.

The K18-human ACE2 transgenic (tg) mouse model, which expresses the human ACE2 receptor, has been widely used to study SARS-CoV-2 because of its susceptibility to infection and ability to produce human-like disease symptoms (20). Several studies have used this model to analyze microbiota in the lungs and gut (21, 22). It is still necessary, however, to comprehensively investigate microbiota changes in the lungs and intestine following infection with different SARS-CoV-2 strains.

By comparing the microbiota in the lungs and intestines of mice infected with SARS-CoV-2 Wuhan and SARS-CoV-2 Omicron, we aimed to uncover strain-specific differences in how SARS-CoV-2 affects the host microbiota. Additionally, we sought to identify microbial signatures associated with each viral strain and explore how differences in microbiota correlate with variations in disease severity and immune response. Understanding these distinctions will provide valuable insights into the pathogenesis of different SARS-CoV-2 variants and inform the development of targeted microbiota-based therapeutic strategies.

## Results

### Differential disease outcomes and microbiota changes induced by SARS-CoV-2 Wuhan and SARS-CoV-2 Omicron in K18-hACE2-tg mice

To evaluate disease outcomes after infection with SARS-CoV-2 Wuhan or SARS-CoV-2 Omicron, we infected K18-hACE2-tg mice with each strain and monitored them for 15 days (**Fig. 1A**). Mice infected with SARS-CoV-2 Wuhan survived for approximately 6 days and showed 100% mortality by the end of the observation period, whereas mice infected with SARS-CoV-2 Omicron showed a mortality rate of only 10% during the observation period (**Fig. 1B**). We investigated viral loads by measuring RNA copies of viral protein N1 (nucleocapsid) in the lung tissues of infected mice (**Fig. 1C**). Despite the difference in mortality, there was no significant difference in viral loads between the two strains at 5 days post infection (dpi); however, the viral load of SARS-CoV-2 Omicron significantly decreased between 5 dpi and 10 dpi. We conducted a histological analysis of lung and tracheal tissues to compare the pathological changes in the respiratory tracts of the infected mice. Compared with non-infected control mice, mice infected with SARS-CoV-2 Wuhan displayed alveolar/interstitial thickening, pulmonary/alveolar hemorrhage, fibrin deposition, and inflammatory cell infiltration in lung tissues, whereas mice infected with SARS-CoV-2 Omicron showed only persistent alveolar/interstitial thickening at 5 dpi and 10 dpi (**Fig. 1D**). The tracheal tissues of the non-infected mice had intact ciliated epithelium, whereas those of the mice infected with SARS-CoV-2 Wuhan displayed severe loss of cilia and partial epithelial detachment, and those of the mice infected with SARS-CoV-2 Omicron displayed reduced tracheal epithelium thickness and partial cilia loss at 5 dpi, with persistent thinning but intact cilia at 10 dpi (**Fig. 1E**). These results demonstrate that infection with SARS-CoV-2 Wuhan caused significant lung and tracheal damage, whereas infection with SARS-CoV-2 Omicron caused relatively mild but persistent respiratory changes over time.

**Figure 1.**
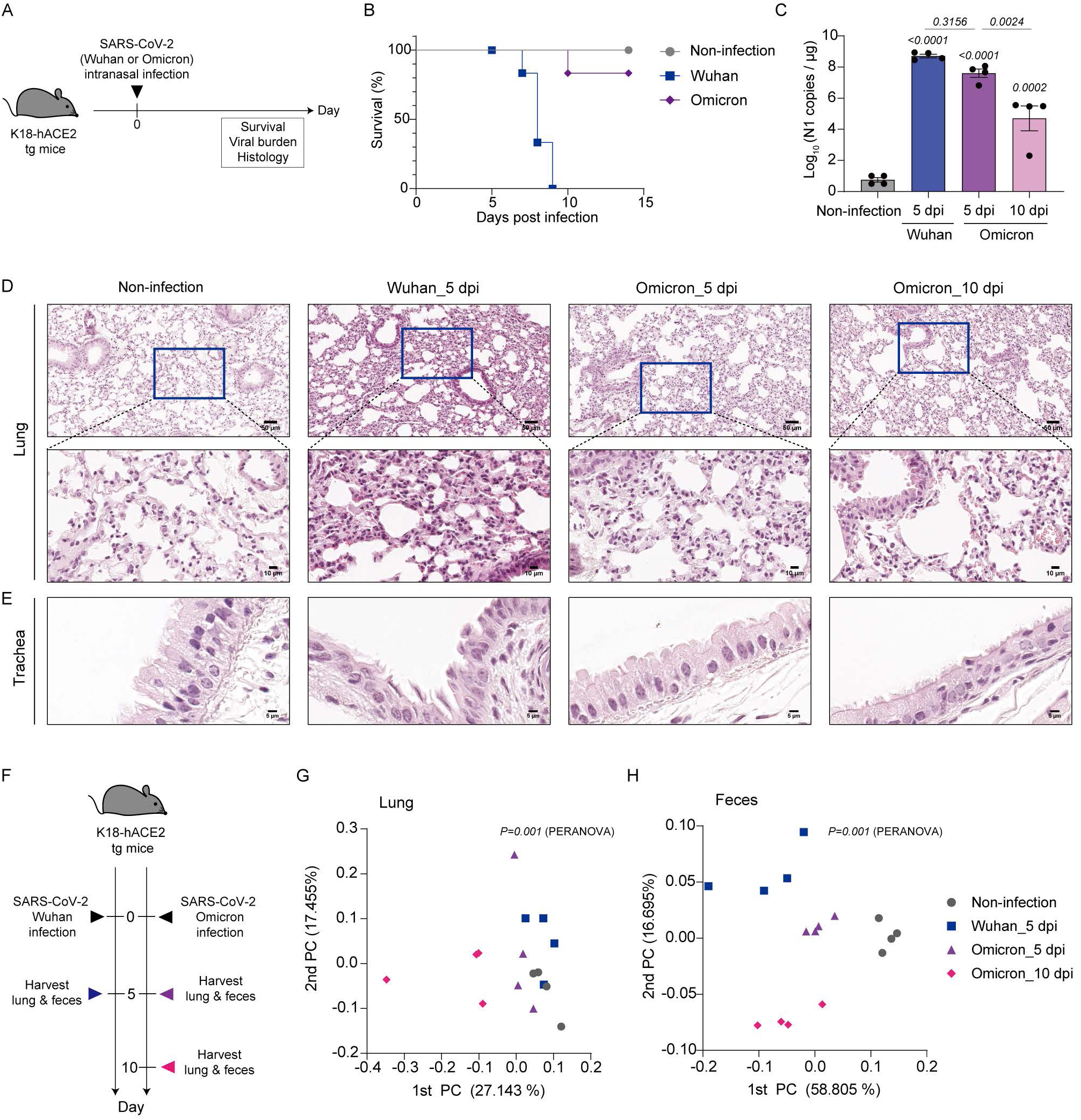
Disease outcomes and beta diversity of lung and intestinal microbiota following infection with SARS-CoV-2 Wuhan or SARS-CoV-2 Omicron. (A) Schematic representation of the timeline of the murine experiments. Six-week-old K18-hACE2-tg mice were intranasally exposed to phosphate-buffered saline (PBS, non-infection), SARS-CoV-2 Wuhan, or SARS-CoV-2 Omicron. (B) Survival rates of the experimental mice. (C) RNA copies of viral N1 in 1 μg of lung tissues total RNA from experimental mice were measured using qPCR. (D, E) Lung and tracheal tissues from experimental mice were harvested at 5 dpi or 10 dpi for histological analysis. Representative images are shown of the lungs (D) and trachea (E). Scale bars in the lung images indicate 50 μm in the first line and 10 μm in the magnified images in the second line. Scale bars in the tracheal images indicate 5 μm. (F) Schematic of the timeline used for sample collection. Six-week-old K18-hACE2-tg mice were intranasally exposed to PBS (non-infection), SARS-CoV-2 Wuhan, or SARS-CoV-2 Omicron. Lung and fecal samples from the non-infection and Wuhan groups were collected at 5 dpi, and samples from the Omicron group were collected at 5 dpi and 10 dpi. (G, H) PCoA plots for lung (G) and fecal (H) samples from the experimental mice. Data in C were analyzed using one-way ANOVA. Data in G and H were analyzed using PERANOVA.

We made two main comparisons regarding the effects of SARS-CoV-2 infections on microbiota: a comparison between SARS-CoV-2 Wuhan and SARS-CoV-2 Omicron at 5 dpi and a comparison between SARS-CoV-2 Omicron at 5 dpi and SARS-CoV-2 Omicron at 10 dpi. For these comparisons, we conducted 16S ribosomal RNA (rRNA) sequencing of lung and fecal DNA samples obtained from infected mice (**Fig. 1F**). Samples from non-infected mice were also collected at 5 dpi. A principal coordinate analysis (PCoA) of the beta diversity of the lung samples showed that the samples collected at 10 dpi from mice infected with SARS-CoV-2 Omicron were distinct from the other samples, which otherwise showed no distinct separation among groups, including the non-infected group (**Fig. 1G**). By contrast, PCoA of the beta diversity of the fecal samples showed that the samples from each group of mice were distinct (**Fig. 1H**). These results demonstrated that infection with different SARS-CoV-2 strains induced distinct changes in the lung and intestinal microbiota of susceptible mice.

### Comparison of lung and intestinal microbiota following infections with SARS-CoV-2 Wuhan and SARS-CoV-2 Omicron

To investigate changes in lung microbiota following infection with SARS-CoV-2 Wuhan or SARS-CoV-2 Omicron, we compared non-infected mice with infected mice at 5 dpi (**Fig. 2A**). First, we considered all infected mice as a single group and compared them with the non-infected mice using the Firmicutes/Bacteroidetes (F/B) ratio and two alpha diversity indices: the ACE index for species richness and the Shannon index for species diversity. Although the values of the diversity indices fluctuated after SARS-CoV-2 infection, there were no significant differences in alpha diversity or F/B ratio between the non-infected and infected mice **(Fig. 2B– 2D)**. We next compared the alpha diversity indices and F/B ratio among the non-infected mice, the mice infected with SARS-CoV-2 Wuhan, and the mice infected with SARS-CoV-2 Omicron (**Fig. 2E–2G**). There were no significant differences in the ACE and Shannon indices among the three groups (**Fig. 2E and 2F**); however, the mice infected with SARS-CoV-2 Omicron exhibited a higher F/B ratio than the mice infected with SARS-CoV-2 Wuhan (**Fig. 2G**), suggesting that the two strains caused different changes in lung microbiota.

**Figure 2.**
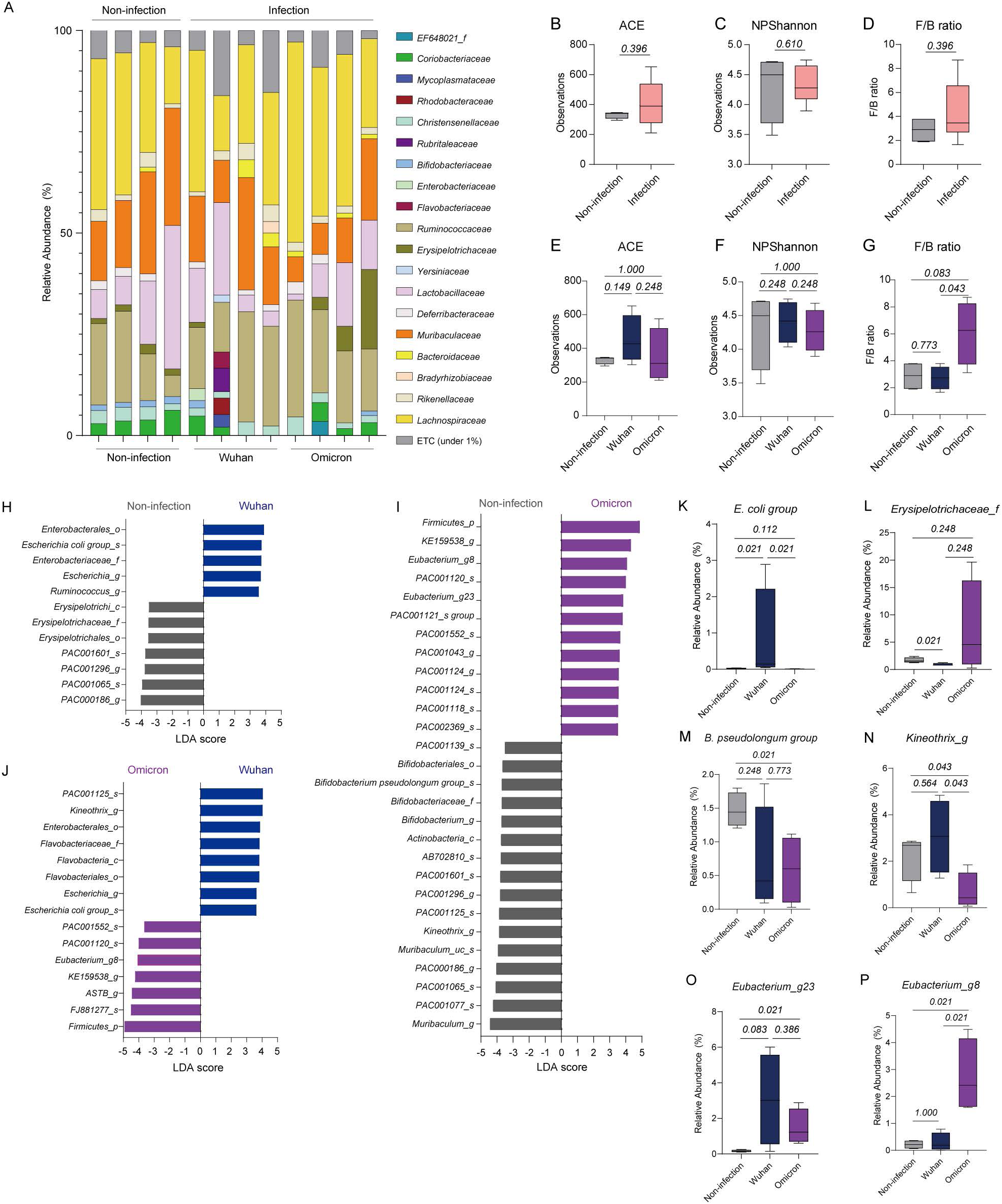
Comparison of lung microbial compositions among non-infected mice and mice infected with SARS-CoV-2 Wuhan or SARS-CoV-2 Omicron. (A) Relative abundances of bacterial families in the lungs of the experimental mice. (B–D) ACE index (B), Shannon index (C), and F/B ratio (D) of the lung microbiota of non-infected mice and mice infected with SARS-CoV-2 (Wuhan or Omicron). (E–G) ACE index (E), Shannon index (F), and F/B ratio (G) of the lung microbiota in non-infected mice, mice infected with SARS-CoV-2 Wuhan, and mice infected with SARS-CoV-2 Omicron. (H–J) Bacteria with LDA scores >3.5 in pairwise comparisons between non-infected mice and mice infected with SARS-CoV-2 Wuhan (H), non-infected mice and mice infected with SARS-CoV-2 Omicron (I), and mice infected with SARS-CoV-2 Wuhan and mice infected with SARS-CoV-2 Omicron (J). (K–P) Relative abundances of *E. coli* group (K)*, Erysieplotrichaceae_f* (L), *B. pseudolongum* group (M), *Kineothrix_g* (N), *Eubacterium_g23* (O), and *Eubacterium_g8* (P) among the lung microbiota of experimental mice. Data in B–G and K–P were analyzed using the Wilcoxon rank-sum test.

To determine which bacteria were specifically affected by SARS-CoV-2 infection, we made pairwise comparisons between groups using linear discriminant analysis (LDA) and sorted the bacteria with LDA scores >3.5 (**Fig. 2H–2J**). Compared with non-infected mice, mice infected with SARS-CoV-2 Wuhan showed enrichment of bacteria related to *Escherichia coli* (*Enterobacterales_o*, *Enterobacteriaceae_f*, *Escherichia_g*) and depletion of *Erysipelotrichaceae_f* and some species of *Muribaculaceae_f* (*PAC000186_g* and *PAC000165_s*) and *Lachnospiraceae_f* (*PAC001296_g* and *PAC001601_s*; **Fig. 2H**). In the comparison between non-infected mice and mice infected with SARS-CoV-2 Omicron, the latter showed enrichment of *Firmicutes_p, Eubacterium_g8*, and *Eubacterium_g23* and depletion of bacteria related to *Bifidobacterium pseudolongum* (*Bifidobacteriales_o*, *Bifidobacteriaceae_f*, *Bifidobacterium_g*), *Muribaculaceae_f* (*Muribaculum_g*, *PAC001077_s, PAC001065_s, PAC000186_g, Muribaculum_uc_s*), and *Lachnospiraceae_f* (*Kineothri_g*, *PAC001125_s, PAC001296_g, PAC001601_s, AB702810_s*; **Fig. 2I**). A direct comparison between mice infected with SARS-CoV-2 Wuhan and mice infected with SARS-CoV-2 Omicron revealed that *Kineothrix_g* and *E. coli* were more prevalent in the former, whereas *Firmicutes_p* and *Eubacterium_g8* were more abundant in the latter (**Fig. 2J**).

Based on these results, we determined that some bacteria were altered among the lung microbiota in specific groups of mice. The abundance of *E. coli* was higher, whereas the abundance of *Erysipelotrichaceae_f* was lower, in the mice infected with SARS-CoV-2 Wuhan than in the non-infected mice and the mice infected with SARS-CoV-2 Omicron (**Fig. 2K and 2L**). The mice infected with SARS-CoV-2 Omicron exhibited a lower abundance of *B. pseudolongum* and *Kineothrix_g* than the non-infected mice (**Fig. 2M and 2N**). In addition, the abundance of *Eubacterium_g23* was higher in the mice infected with SARS-CoV-2 Wuhan or SARS-CoV-2 Omicron compared with that in the non-infected mice, but the abundance of *Eubacterium_g8* was elevated only in the mice infected with SARS-CoV-2 Omicron (**Fig. 2O and 2P**). These results indicated that the lung microbiota changed specifically in response to infection with each SARS-CoV-2 strain, suggesting that the two strains differentially induced changes in the lung microbiome composition.

We next conducted a similar analysis using fecal samples from K18-hACE2 tg mice infected with SARS-CoV-2 (**Fig. 3A**). We first compared the alpha diversity indices and F/B ratio between mice infected with SARS-CoV-2 Wuhan or SARS-CoV-2 Omicron and non-infected mice (**Fig. 3B–3D**). The ACE index values did not significantly change following infection (**Fig. 3B**); however, the Shannon index values were higher in the SARS-CoV-2–infected mice than in the non-infected mice (**Fig. 3C**), suggesting that SARS-CoV-2 infection decreased species diversity, but not richness, in the intestine. Additionally, the F/B ratio was higher in the SARS-CoV-2–infected mice than in the non-infected mice (**Fig. 3D**). When we compared these values between the mice infected with SARS-CoV-2 Wuhan and those infected with SARS-CoV-2 Omicron, there was no significant difference in ACE and Shannon indices between the two groups (**Fig. 3E and 3F**); however, the F/B ratio was higher in the mice infected with SARS-CoV-2 Wuhan than in the mice infected with SARS-CoV-2 Omicron or the non-infected mice (**Fig. 3G**).

**Figure 3.**
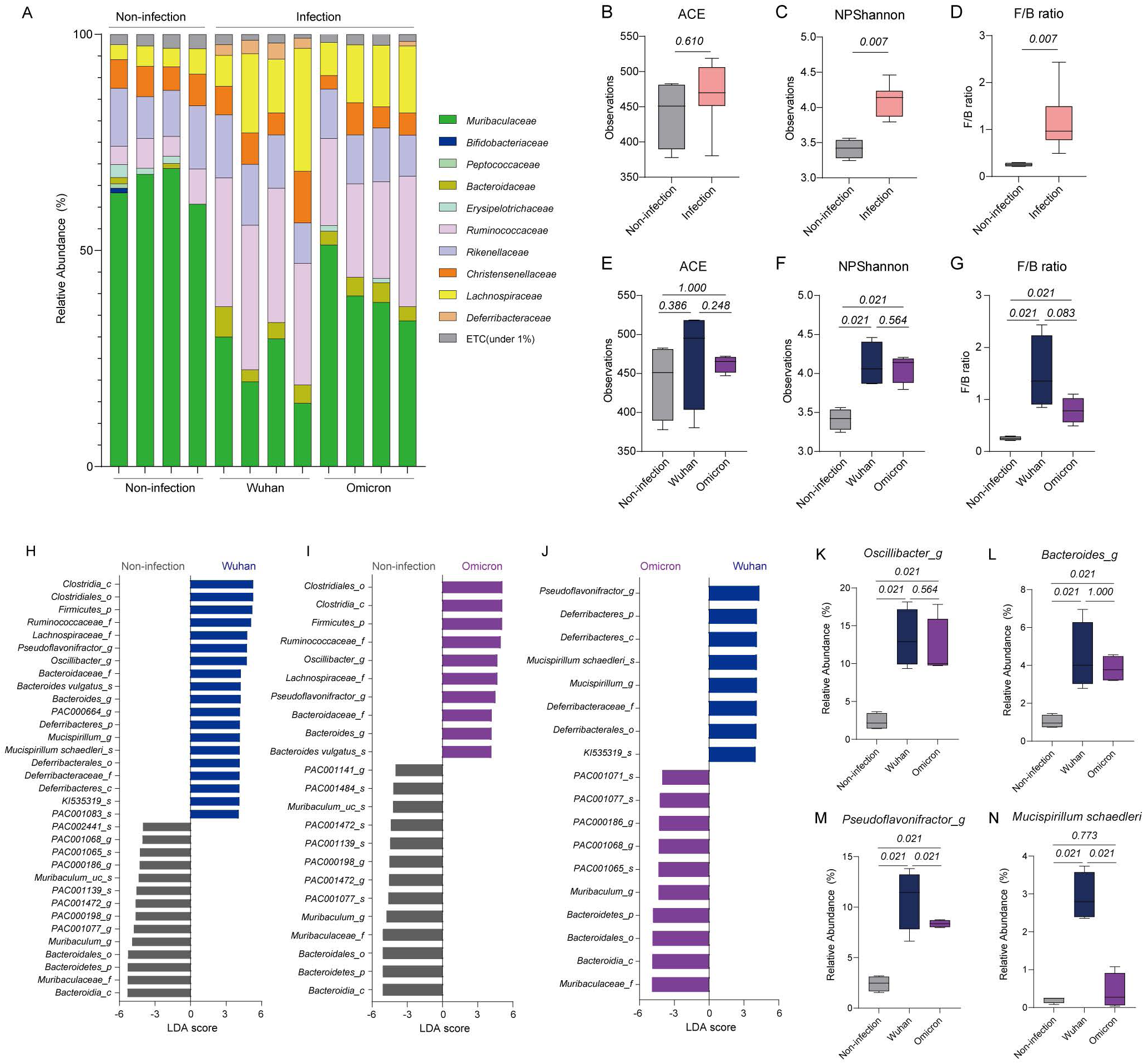
Comparison of intestinal microbial compositions among the non-infection, Wuhan and Omicron groups. (A) Relative abundances of bacterial families in the intestines of the experimental mice. (B–D) ACE index (B), Shannon index (C), and F/B ratio (D) of the intestinal microbiota of non-infected mice and mice infected with SARS-CoV-2 (Wuhan or Omicron). (E–G) ACE index (E), Shannon index (F), and F/B ratio (G) of the intestinal microbiota of non-infected mice, mice infected with SARS-CoV-2 Wuhan, and mice infected with SARS-CoV-2 Omicron. (H-J) Bacteria with LDA scores >4 in pairwise comparisons between non-infected mice and mice infected with SARS-CoV-2 Wuhan (H), non-infected mice and mice infected with SARS-CoV-2 Omicron (I), and mice infected with SARS-CoV-2 Wuhan and mice infected with SARS-CoV-2 Omicron (J). (K–N) Relative abundances of *Oscillibacter_g* (K)*, Bacteroides_g* (L), *Pseudoflavonifractor_g* (M), and *Mucispirillum schaedleri* (N) among the intestinal microbiota of experimental mice. Data in B–G and K–N were analyzed using the Wilcoxon rank-sum test.

Next, we conducted a comparison among the non-infected mice, the mice infected with SARS-CoV-2 Wuhan, and the mice infected with SARS-CoV-2 Omicron using the LDA method and subsequently sorted the bacteria with LDA scores >4 (**Fig. 3H–3J**). Compared with the non-infected mice, the mice infected with SARS-CoV-2 Wuhan exhibited a higher abundance of genera including *Pseudoflavonifractor*, *Oscillibacter*, *Bacteroides*, and *Mucispirillum* (**Fig. 3H**). Similarly, the mice infected with SARS-CoV-2 Omicron showed enrichment of genera such as *Oscillibacter*, *Pseudoflavonifractor*, and *Bacteroides*, but not *Mucispirillum*, compared with the non-infected mice (**Fig. 3I**). The mice infected with SARS-CoV-2 Wuhan exhibited enrichment of bacteria related to *Mucispirillum schaedleri* (*Mucispirillum_g, Deferribacteraceae_f, Deferribacterralse_o, Deferribacteres_c,* and *Deferribacteres_p)* and *Pseudoflavonifractor_g* compared with the mice infected with SARS-CoV-2 Omicron (**Fig. 3J**).

After conducting these comparisons, we confirmed the relative abundances of *Oscillibacter_g, Bacteroides_g, Pseudoflavonifrator_g*, and *Muscispirillum schaedleri* in the non-infected mice and the mice infected with each SARS-CoV-2 strain (**Fig. 3K–3N**). The abundances of *Oscillibacter_g* and *Bacteroides_g* notably increased in mice infected with either SARS-CoV-2 strain compared with those in the non-infected mice; however, no substantial differences were observed between the mice infected with SARS-CoV-2 Wuhan and the mice infected with SARS-CoV-2 Omicron (**Fig. 3K and 3L**). Mice infected with either SARS-CoV-2 strain exhibited a higher abundance of *Pseudoflavonifractor_g* than the non-infected mice, and the abundance of *Pseudoflavonifractor_g* in the mice infected with SARS-CoV-2 Wuhan was significantly higher than that in the mice infected with SARS-CoV-2 Omicron (**Fig. 3M**). *M. schaedleri* was significantly more abundant in the mice infected with SARS-CoV-2 Wuhan than in the non-infected mice or the mice infected with SARS-CoV-2 Omicron (**Fig. 3N**). These results demonstrate both common and specific changes in the intestinal microbiota after infection with SARS-CoV-2 Wuhan or SARS-CoV-2 Omicron.

### Alterations in lung and intestinal microbiota over time following SARS-CoV-2 Omicron infection

Although infection with SARS-CoV-2 Omicron was not life-threatening, it caused damage to the lung and tracheal barriers in K18-hACE-2 tg mice and produced a viral load similar to that of life-threatening SARS-CoV-2 Wuhan infection, which persisted for up to 10 days after infection (**Fig. 1**). Therefore, we investigated the changes in lung microbiota at 5 days and 10 days after SARS-CoV-2 Omicron infection (**Fig. 4A**). We also compared the alpha diversity indices and F/B ratios between non-infected mice and mice infected with SARS-CoV-2 Omicron at 5 dpi and 10 dpi (**Fig. 4B–4D**). The ACE index was significantly decreased at 10 dpi compared with that in the non-infected mice, but the Shannon index did not change over the course of the infection (**Fig. 4B and 4C**). Furthermore, the F/B ratio, which was increased at 5 dpi, remained constant at 10 dpi (**Fig. 4D**).

**Figure 4.**
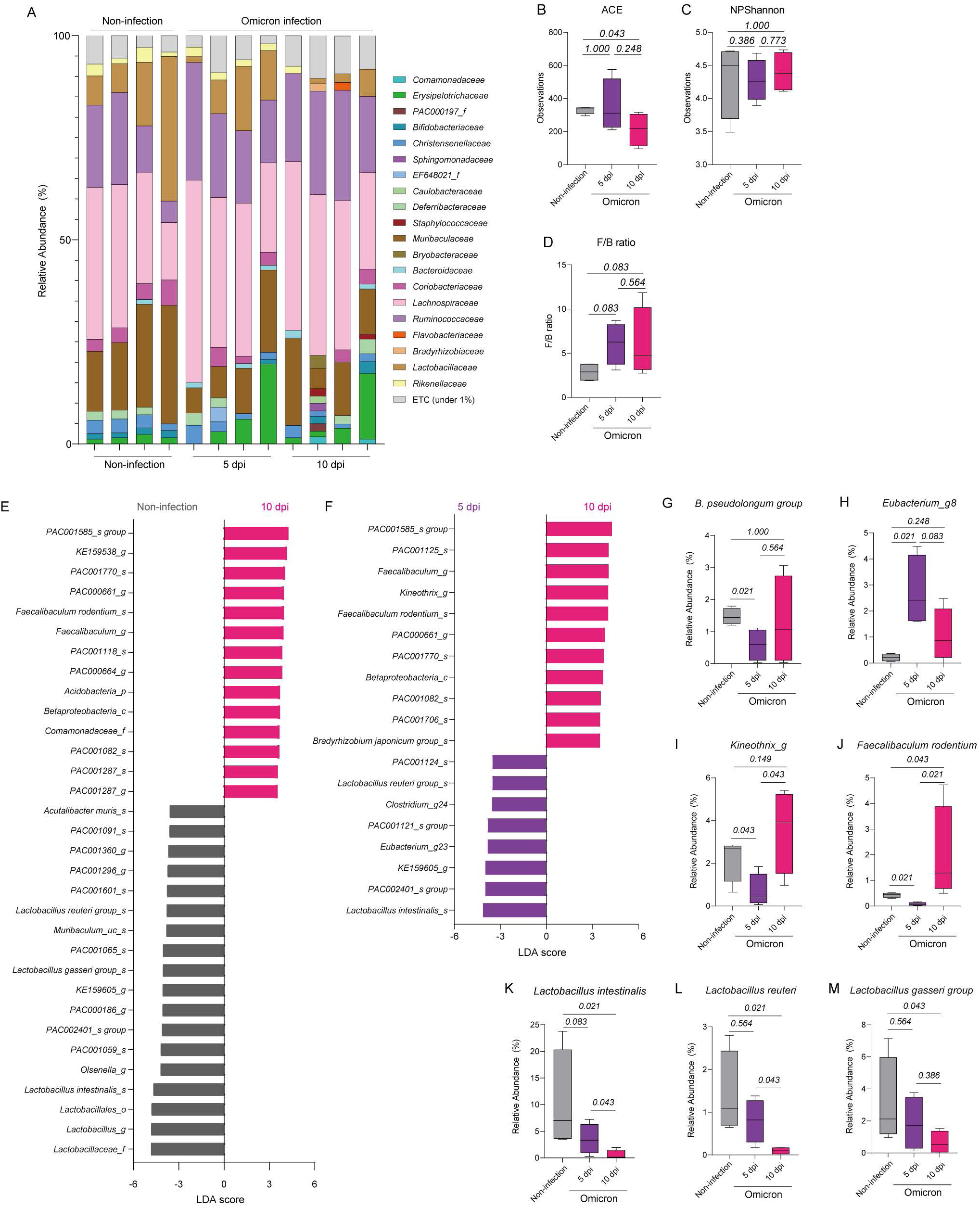
Comparison of lung microbiota over time following infection with SARS-CoV-2 Omicron. (A) Relative abundances of bacterial families among lung microbiota of non-infected mice and mice infected with SARS-CoV-2 Omicron at 5 dpi and 10 dpi. (B–D) ACE index (B), Shannon index (C), and F/B ratio (D) of the lung microbiota in non-infected mice and mice infected with SARS-CoV-2 Omicron at 5 dpi and 10 dpi. (E, F) Bacteria with LDA scores >3.5 in pairwise comparisons of lung microbiota between non-infected mice and mice infected with SARS-CoV-2 Omicron (F) and mice infected with SARS-CoV-2 Omicron at 5 dpi and 10 dpi (F). (G–M) Relative abundances of *B. pseudolongum* (G)*, Eubacterium_g8* (H), *Kineothrix_g* (I), *Faecalibculum rodentium* (J), *Lactobacillus intestinalis* (K), *Lactobacillus reuteri* (L), and *Lactobacillus gasseri* (M) among lung microbiota of experimental mice. Data in B–D and G–M were analyzed using the Wilcoxon rank-sum test.

We next determined which bacteria were specifically enriched among the lung microbiota at 10 dpi in the mice infected with SARS-CoV-2 Omicron using the LDA method (**Fig. 4E and 4F**). Compared with non-infected mice, mice infected with SARS-CoV-2 Omicron showed enrichment of *PAC001585_s* and *PAC000661_g* of *Oscillospiraceae_f*; *KE159538_g, PAC001770_s, PAC001118_s, PAC000664_g, PAC001082_s, PAC001287_s,* and *PAC001287_g* of *Lachnospiraceae_f*; *Faecalibaculum rodentium*; *Acidobactera_p*; *Betaproteobacteria_c*; and *Comamonadaceae_f*, along with depletion of bacteria related to the *Lactobacillus* genus (*Lactobacillaceae_f, Lactobacillacles_o, Lactobacillus intestinalis, Lactobacillus gasseri* group, and *Lactobacillus reuteri* group; **Fig. 4E**). The lung microbiota of mice infected with SARS-CoV-2 Omicron also showed enrichment of *F. rodentium* and depletion of *L. intestinalis* and *L. reuteri* at 10 dpi compared with 5 dpi (**Fig. 4F**).

Next, we confirmed the relative abundances of *B. pseudolongum* and *Eubacterium_g8,* which differed between the non-infected mice and the mice infected with SARS-CoV-2 Omicron at 5 dpi (**Fig. 4G and 4H**). Unlike the infected mice at 5 dpi, the infected mice at 10 dpi exhibited abundances of these bacteria similar to those in the non-infected group (**Fig. 4G and 4H**). In addition, the abundance of *Kineothrix_g* was decreased at 5 dpi compared with that in non-infected mice and subsequently increased from 5 dpi to 10 dpi; however, the abundance at 10 dpi was not significantly different than that in the non-infected mice (**Fig. 4I**). The abundance of *F. rodentium* was similarly decreased at 5 dpi but was significantly elevated at 10 dpi compared with that in the non-infected mice (**Fig. 4J**). We also observed that the abundance of several *Lactobacillus* species was decreased at 10 dpi (**Fig. 4K–4M**). These findings imply that the lung microbiota undergoes persistent modifications following SARS-CoV-2 infection.

We next analyzed the microbial composition using 16S rRNA sequencing data from fecal samples of non-infected mice and mice infected with SARS-CoV-2 at 5 dpi and 10 dpi (**Fig. 5A**). There were no significant changes in ACE index among the three groups (**Fig. 5B**); however, the Shannon index and F/B ratios of the infected mice at 10 dpi were higher than those of the non-infected mice and similar to those of the infected mice at 5 dpi (**Fig. 5C and 5D**). These results demonstrated that the species richness, species diversity, and F/B ratio of the intestinal microbiota did not significantly change from 5 dpi to 10 dpi.

**Figure 5.**
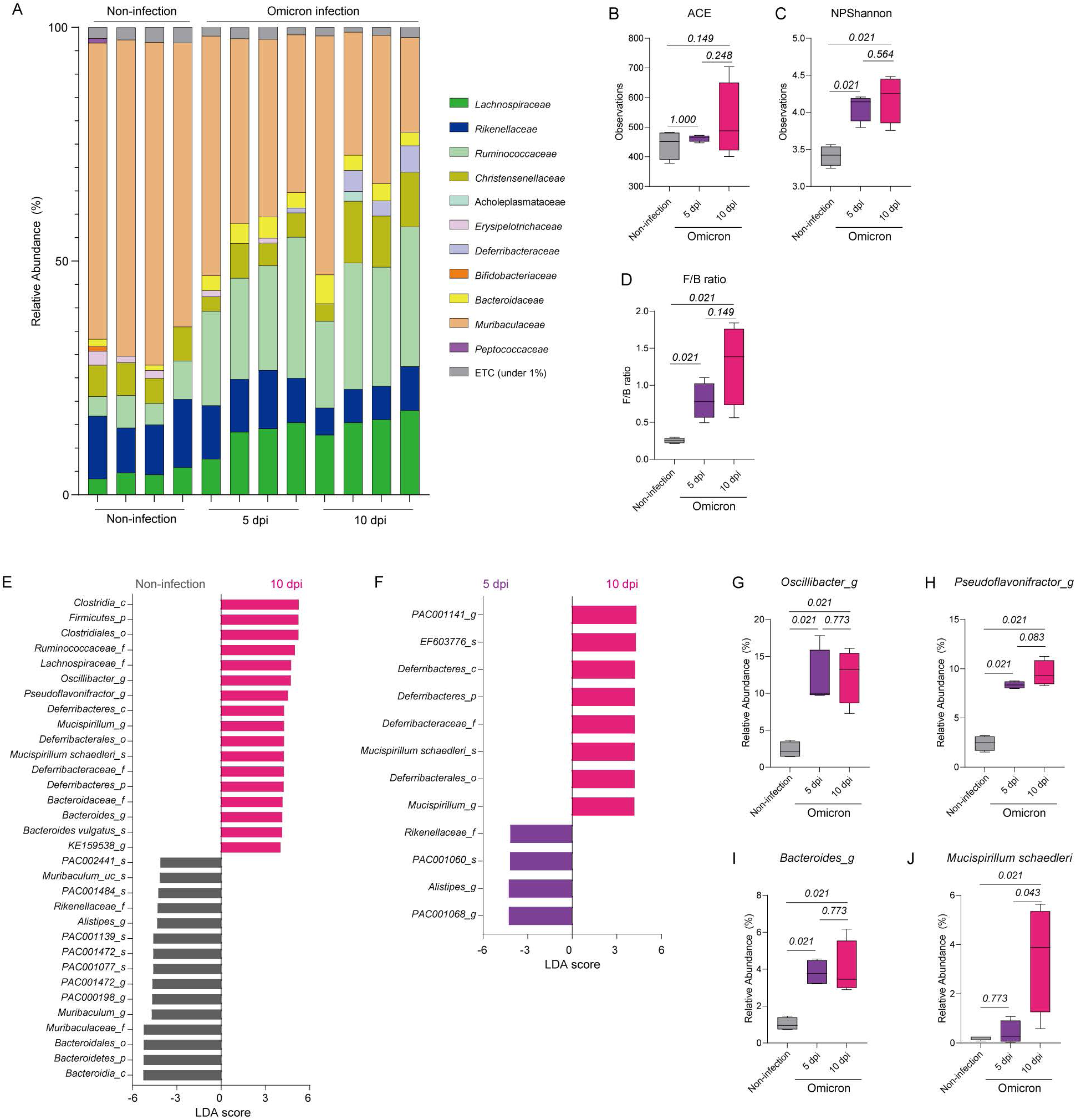
Comparison of intestinal microbiota over time following infection with SARS-CoV-2 Omicron. (A) Relative abundances of bacterial families among intestinal microbiota in non-infected mice and mice infected with SARS-CoV-2 Omicron at 5 dpi and 10 dpi. (B–D) ACE index (B), Shannon index (C), and F/B ratio (D) of the intestinal microbiota of non-infection mice and mice infected with SARS-CoV-2 Omicron at 5 dpi and 10 dpi. (E, F) Bacteria with LDA scores >4.0 in pairwise comparisons between non-infected mice and mice infected with SARS-CoV-2 at 10 dpi (E) and mice infected with SARS-CoV-2 at 5 dpi and 10 dpi (F). (G–J) Relative abundances of *Oscillibacter_g* (G)*, Pseudoflavonifrator_g* (H), *Bacteroides_g* (I) and *Mucispirillum schaedleri* (J) among intestinal microbiota of experimental mice. Data in B–D and G–J were analyzed using the Wilcoxon rank-sum test.

Using the LDA method, we determined which intestinal bacteria were enriched at 10 dpi compared with 5 dpi and non-infected mice (**Fig. 5E and 5F**). Similar to the microbiota at 5 dpi, the microbiota at 10 dpi exhibited enrichment of bacteria including *Ruminococcaceae_f, Lachnospiraceae_f, Oscillibacter_g*, *Pseudoflavonifractor_g,* and *Bacteroides_g* compared with the microbiota of non-infected mice (**Fig. 5E**). In addition, *M. schaedleri*, which was enriched in the intestinal microbiota of mice infected with SARS-CoV-2 Wuhan, was also enriched at 10 dpi compared with both 5 dpi and non-infected mice (**Fig. 5E and 5F**). We confirmed the relative abundances of *Oscillibacter_g*, *Pseudoflavonifractor_g, Bacteroides_g,* and *M. schaedleri* in the three groups (**Fig. 5G–5J**). The abundances of *Oscillibacter_g*, *Pseudoflavonifractor_g,* and *Bacteroides_g* were increased at 5 dpi and remained high until 10 dpi (**Fig. 5G–5I**); however, the abundance of *M. schaedleri* was significantly increased only at 10 dpi compared with that in non-infected mice (**Fig. 5J**). These results suggest that while the alterations of intestinal microbiota due to SARS-CoV-2 Omicron infection were maintained, additional changes gradually occurred over the course of infection.

## Discussion

We investigated the changes in lung and intestinal microbiota in K18-hACE2-tg mice infected with SARS-CoV-2 Wuhan or SARS-CoV-2 Omicron. SARS-CoV-2 Wuhan was 100% lethal, whereas SARS-CoV-2 Omicron had lower lethality. Therefore, we compared the microbiota of mice infected with SARS-CoV-2 Omicron at 5 dpi, when the viral load was similar to that of mice infected with SARS-CoV-2 Wuhan, with that at 10 dpi, when the viral load had decreased but remained detectable. Our comparisons demonstrated differences in microbiota between mice infected with each strain and showed that changes in microbiota occurred over time in mice infected with SARS-CoV-2 Omicron.

Analysis of the lung microbiota revealed no significant differences in the ACE and Shannon indices between mice infected with each strain. A previous study reported that infection with a high dose (1×10^5^ PFU) rather than a low dose (1×10^4^ PFU) of SARS-CoV-2 Wuhan induced an increase in the F/B ratio (21). Our experimental dose (5×10^4^ PFU) of SARS-CoV-2 Wuhan did not induce a change in the F/B ratio; however, the same dose of SARS-CoV-2 Omicron did increase the F/B ratio. In addition, our analysis demonstrated that more bacteria, such as the *B. pseudolongum* group*, Keneothrix_g,* and *Eubacterium_g8,* were specifically altered by SARS-CoV-2 Omicron infection than by SARS-CoV-2 Wuhan infection. These results imply that compared with SARS-CoV-2 Wuhan, SARS-CoV-2 Omicron may be more likely to induce dysbiosis of the lung microbiome when an equal amount of virus is introduced, regardless of lethality.

We found that SARS-CoV-2 Wuhan infection led to increased *E. coli* abundance and decreased *Erysipelotrichaceae_f* abundance among the lung microbiota and also caused significant damage to the barriers of the lungs and trachea. Some bacterial infections occur during viral pneumonia following SARS-CoV-2 infection and are likely to contribute to mortality (23, 24). *E. coli* is one of the pathogens discovered in COVID-19 patients (25). In addition, one study reported that the abundance of *Erysipelotrichaceae_f* was negatively associated with the concentration of IL-4 in the lungs (26). In the K18-hACE2-tg mouse model, a cytokine storm including IL-6, IL-17, and IL-4 occurred in the lungs following SARS-CoV-2 Wuhan infection (20). Another study showed that patients infected with SARS-CoV-2 Wuhan had higher levels of cytokines, including IL-4, in plasma serum than patients infected with SARS-CoV-2 Omicron (27). Therefore, it is suggested that barrier disruption and cytokine storm caused by SARS-CoV-2 Wuhan infection can lead to specific changes in the abundances of *E. coli* and *Erysipelotrichaceae_f* among the lung microbiota.

Infection with either SARS-CoV-2 strain induced changes in the Shannon index but not in the ACE index of the intestinal microbiota. Although the F/B ratio of the lung microbiota remained unchanged after infection with SARS-CoV-2 Wuhan, the F/B ratio of the intestinal microbiota increased following the same infection. In addition, we found that *M. schaedleri* was specifically increased in the intestinal microbiome after SARS-CoV-2 Wuhan infection. This bacterium has been reported to be not only an antagonist of Salmonella infection but also an indicator of colitis in dextran sulfate sodium-induced and immune-deficient murine models (28–31). These findings suggest that the Wuhan strain may induce intestinal inflammation more rapidly than the Omicron strain in our murine model.

We observed that considerable viral loads in lung tissues persisted 10 days after initial SARS-CoV-2 Omicron infection. In the comparison of lung microbiota over time following SARS-CoV-2 Omicron infection, the abundances of some bacteria, such as *B. pseudolongum, Eubacgerium_g8,* and *Kineothrix_g*, changed at 5 dpi but became similar to those in non-infected mice at 10 dpi. However, we also found that the abundance of *Faecalibaculum rodentium* increased, particularly at 10 dpi. *F. rodentium* and its human homolog, *Holdemanella biformis*, have been reported to suppress murine colitis and protect against intestinal tumor development (32, 33). One study using Mendelian randomization analysis reported a causal association between intestinal *Holdemanella* and allergic asthma (34); however, the role of this bacterium in the lungs remains unclear. We also observed a decrease in *Lactobacillus spp.* abundance at 10 dpi following SARS-CoV-2 Omicron infection. A previous study demonstrated enrichment of *Lactobacillus* in the lung microbiota of COVID-19 patients compared with healthy controls (35). However, some studies have reported that intranasal delivery of *Lactobacillus spp.* protects against respiratory viral and bacterial infections (36, 37). Therefore, it remains unclear how the bacteria that were altered at 10 dpi behave in the lungs, and further studies are necessary to determine the association between SARS-CoV-2 infection and these bacteria.

Although the alterations of lung microbiota observed at 5 dpi did not persist until 10 dpi, the increased Shannon index and F/B ratio of the intestinal microbiota observed at 5 dpi were still present at 10 dpi. In addition, the abundances of the bacteria that were altered at 5 dpi, such as *Oscillibacter_g, Pseudoflavonifractor_g,* and *Bacteroides_g*, were not significantly altered at 10 dpi, suggesting that the alteration of intestinal microbiota remained constant following SARS-CoV-2 Omicron infection. *Oscillibacter_g* was discovered among human gut microbiota associated with Crohn’s disease and mesenteric fat inflammation (38, 39). In addition, among the *Bacteroides_g, B. vulgatus* was enriched in the intestinal microbiome at 10 dpi after SARS-CoV-2 Omicron infection (**Fig. 5E**). A previous study showed that patients with long COVID symptoms exhibited changes in the gut microbiome, including higher levels of *B. vulgatus* (40). Moreover, *M. schaedleri*, which was enriched in the intestinal microbiota of mice infected with SARS-CoV-2 Wuhan at 5 dpi, was particularly enriched in mice infected with SARS-CoV-2 Omicron at 10 dpi. These results suggest that although SARS-CoV-2 Omicron infection may not induce intestinal inflammation as rapidly as SARS-CoV-2 Wuhan infection, it might still affect intestinal inflammation at a later stage.

This study provides a comprehensive comparison of the pathological and microbiological effects of the Wuhan and Omicron variants of SARS-CoV-2 in a susceptible mouse model. By elucidating the differential outcomes and microbial changes associated with these variants, we aimed to enhance our understanding of COVID-19 pathogenesis and inform future therapeutic and preventive measures against emerging variants. Further studies with larger sample sizes and considerations for cage effects are needed to better understand the role of microbiome changes in SARS-CoV-2 infection, particularly in the intestinal and respiratory tracts. In addition, further research is needed to investigate the continuous changes in microbiota following infection with non-lethal strains of SARS-CoV-2 through prolonged observation periods longer than 10 days.

## Materials and Methods

### Ethics statement

The animal study was conducted in strict accordance with the recommendations of the Guide for the Care and Use of Laboratory Animals (Protocol No. 8A-M6, 2023-8A-07-03) from Korea Research Institute of Chemical Technology (KRICT). Infection was performed in a BL-3 containment facility at KRICT.

### Cell lines and viruses

Vero cells (CCL-81) were purchased from the American Type Culture Collection (ATCC, Manassas, VA, USA) and maintained at 37°C with 5% CO_2_ in Dulbecco’s modified Eagle’s medium (DMEM; Gibco, Waltham, MA, USA) supplemented with 10% heat-inactivated fetal bovine serum (FBS; Gibco) and 1% penicillin-streptomycin (Gibco). SARS-CoV-2 Wuhan-Hu-1 (hCoV/Korea/KCDC03/2020) and SARS-CoV-2 Omicron B.1.1.529 (hCoV-19/Korea/KDCA447321/2021) were provided by the National Culture Collection for Pathogens (Cheongju-si, Korea). Viral stock propagation and titers were measured by plaque assay on Vero cells.

### Mice

Animal studies were conducted in accordance with the guidelines outlined in the Guide for the Care and Use of Laboratory Animals of the Korea Advanced Institute of Science and Technology (KAIST) and KRICT. Male K18-hACE2 transgenic mice (strain #034860: B6.Cg-Tg(K18-ACE-2)2Prlmn/J) aged 6–8 weeks were obtained from The Jackson Laboratory and housed under specific pathogen-free conditions at the KAIST Laboratory Animal Resource Center. The infection study was conducted at the KRICT-BL3 facility. Intranasal virus inoculation was performed using 5×10^4^ PFU of SARS-CoV-2 Wuhan or SARS-CoV-2 Omicron. To minimize animal suffering, anesthesia was induced and maintained with isoflurane during virus inoculation and minor treatments.

### Bacterial DNA isolation from mouse stool

For gut microbiome analysis, fecal samples were collected at 5 dpi and 10 dpi and immediately stored at -80 °C for total DNA extraction. DNA extraction was performed using the QIAamp DNA Stool Mini Kit (Qiagen, Hilden, Germany) according to the manufacturer’s instructions. Briefly, a 200 mg fecal sample from each mouse was homogenized using a vortex mixer. The sample was then lysed at 95°C for 5 min, and the debris was pelleted by centrifugation at 12,000 rpm for 1 min. The supernatant was mixed with proteinase K in a new tube, and 200 μL Buffer AL was subsequently added and mixed by vortexing. The mixture was incubated at 70°C for 10 min, and 200 μL molecular-grade pure ethanol was then added to the lysate. The lysate was applied to a QIAamp spin column and centrifuged at 12,000 rpm for 1 min. The column was washed twice with washing buffers (Buffers AW1 and AW2), and 100 μL elution buffer (Buffer ATE) was finally added. The column was centrifuged at maximum speed for 1 min to extract the DNA for sequencing and analysis.

### Bacterial DNA isolation from mouse lungs

Simultaneously with the gut microbiome analysis, total lung tissues were collected at 5 dpi and 10 dpi. The infected mice were euthanized, and approximately 25 mg lung tissue was lysed in 180 µL enzymatic lysis buffer (10 mM Tris-HCl, pH 8.0; 2 mM EDTA, pH 8.0; and 1.2% Triton X-100) containing lysozyme (20 mg/mL). The lysate was incubated at 37°C for 30 min and then at 56°C for another 30 min after addition of proteinase K (0.2 mg/mL). Subsequently, DNA extraction was performed using the DNeasy Blood & Tissue Kit (Qiagen) following the manufacturer’s instructions.

### Viral load quantification

Total RNA was extracted from the right lung using the Qiagen RNeasy Mini kit (Qiagen, Germantown, MD, USA) following the manufacturer’s protocol. Purified total RNA was used for quantification of viral RNA with one-step PrimeScript III RT-qPCR mix (RR600A, Takara, Kyoto, Japan) and a CFX96 Real-Time PCR system (Bio-Rad, Hercules, CA, USA). The viral nucleoprotein (NP) RNA was detected and analyzed using a 2019-nCoV-N1 probe (10006770, Integrated DNA Technologies, Coralville, IA, USA).

### Lung sectioning and histology

To examine the pathological changes in the respiratory tracts of infected mice, mice were euthanized by CO_2_ inhalation. The left lung lobes and trachea were then fixed in 10% neutral-buffered formalin. The fixed tissues were embedded in paraffin and sectioned at a thickness of 5 µm for hematoxylin and eosin (H&E; BBC Biochemical, Mount Vernon, WA, USA) staining. The lung and tracheal tissue sections were examined using a digital slide scanner (3D Histech, Budapest, Hungary), and the severity of tissue damage was assessed across all lung lobes and the trachea.

### 16S rRNA sequencing

Bacterial DNA amplification was conducted using PCR targeting the V3-V4 regions of the 16S rRNA gene under the following conditions: initial denaturation at 95°C for 3 min, followed by 30 cycles at 95°C for 30 s, 55°C for 30 s, and 72°C for 30 s, and final elongation at 72°C for 5 min. Low-quality reads with average quality scores <25 were removed using Trimmomatic v.0.32. Quality-controlled paired-end sequence data were merged using VSERACH version 2.13.4, and primer sequences were trimmed using an alignment algorithm. Non-specific amplicons that did not encode 16S rRNA were identified using the HMMER software package ver.3.2.1. Unique reads were extracted, and redundant reads were clustered with unique reads using the derep_fulllength command of VSEARCH. Taxonomic assignment was performed using the EzBioCloud 16S rRNA database with precise pairwise alignment (41). Chimeric reads were filtered using the UCHIME algorithm and non-chimeric 16S rRNA database from EzBioCloud, removing reads with <97% similarity. After chimeric filtering, reads that could not be identified at the species level with <97% similarity in the EzBioCloud rRNA database were compiled, and *de novo* clustering was performed using the cluster_fast command to generate additional operational taxonomic units (OTUs). OTUs consisting of single reads (singletons) were excluded from further analysis.

Secondary analyses, including diversity calculations and biomarker discovery, were conducted using in-house programs from CJ Bioscience Inc. (Seoul, South Korea). The alpha diversity ACE and Shannon indices were estimated, and beta diversity distances were calculated using the Jensen-Shannon method to visualize sample differences. Taxonomic and functional biomarkers were identified using statistical comparison algorithms (LDA Effect Size, LefSe) with the functional profiles predicted by PICRUSt. All analyses were performed using EzBioCloud 16S-based MTP and CJ Bioscience’s bioinformatics cloud platform.

### Statistical analysis

Viral loads were compared using the one-way ANOVA function of GraphPad Prism version 10 (GraphPad Software, Inc., San Diego, CA, USA). The Wilcoxon rank-sum test in EzBioCloud (CJ Bioscience) was used to compare values, including ACE index, Shannon index, F/B ratio, and abundance, between groups. PCoA data were analyzed using PERMANOVA. Data are presented as Min to Max using GraphPad Prism version 10.

## Acknowledgements

The authors thank the members of the Laboratory of Host Defenses for their helpful discussions. This work was supported by National Research Foundation of Korea grants (NRF-2021M3A9H3015688, NRF-2021M3A9D3026428, NRF-2023R1A2C3003825, and RS-2024-00342560). This work was also supported by Korea Research Institute of Chemical Technology grant (K-GRC GO! KRICT project BSF24-113).

